# Germination timing under climate change: warmer springs favor early germination of range-wide cork oak populations

**DOI:** 10.1101/2023.01.13.523208

**Authors:** Marta Benito Garzón, Fany Baillou, Filipe Costa e Silva, Carla Faria, Maurizio Marchi, Bouthenia Stiti, Giovanni Giuseppe Vendramin, Natalia Vizcaíno-Palomar

**Author notes:** Corresponding author: Marta Benito Garzón.

## Abstract

Climate change is favoring the northward shift of Mediterranean species which are expanding their ranges at their leading edges, becoming natural candidates for increasing forest biodiversity in these regions. However, current knowledge on tree populations’ responses to climate change is mostly based on adult trees, even if tree early developmental stages are far more sensitive to climate and tightly linked to fitness. To fill this knowledge gap, we investigated the potential adaptation of cork oak range-wide populations to increasing spring temperature in germination and post-germination traits. We sowed 701 acorns from 11 populations at 15, 20 and 25°C, monitored germination daily and measured post-germination traits. We model germination timing through Cox’s proportional-hazards models, assess populations’ adaptation to spring temperature transfer distances and quantify the effect of acorn mass and storage duration on all considered traits with fixed-effects models. We predict germination and post-germination climate niches under current and RCP 8.5 2080 scenarios. Large differences in germination timing are due to both the population origin and temperature treatment; germination and survival rates showed a sub-optimality towards warmer-than-origin temperatures and heavier acorns produced faster growing seedlings. The timing of germination is the early stage trait most affected by increasing spring temperatures, with germination in 2080 predicted to be 12 days earlier than to date in the northern part of the species’ range. Warmer spring temperatures will significantly accelerate the germination of other recalcitrant Mediterranean species, which could alter seedlings developmental environment and ultimately populations’ regeneration and species composition. As such, germination timing should receive more attention by scientists and stakeholders, and should be included in forest vulnerability assessments and assisted migration programs aiming at long-term forest regeneration to adapt forests to climate change.

## 1. Introduction

Climate change is disrupting biological cycles, altering ecosystems’ functioning and changing forests communities at an unprecedented pace. Tree populations can survive under climate change by moving towards more favorable conditions (Chen et al., 2011), or persisting *in-situ* by evolutionary processes, including genetic adaptation and phenotypic plasticity (Valladares et al., 2014). In trees, plasticity can promote a fast adjustment to new conditions as those expected with climate change (Valladares et al., 2014) whereas gene flow from core to marginal populations may promote adaptation lags as those observed at population margins (Fréjaville et al., 2020; Rehfeldt et al., 2002). Current knowledge on tree populations’ responses to climate change is mostly based on adult trees, even though early developmental stages are far more sensitive to climate than adult ones and tightly linked to fitness (Verdú and Traveset, 2005), and phenotypic expression changes through developmental stages (Vizcaíno-Palomar et al., 2020). This adult-based knowledge gives only a partial view of forest vulnerability to climate change (Saatkamp et al., 2019).

The very few approaches that investigate range-wide trees’ early developmental stages highlight that germination often shows a narrower climatic niche than adult traits (Walck et al., 2011). This niche difference is the consequence of natural selection and primary environmental filtering, potentially constraining species ranges and promoting narrower niches depending on populations germination sensitivity to new climatic conditions (Donohue et al., 2010). The germination niche is triggered by the environmental cues and genetic determinism of seeds that change across species ranges (Jiménez-Alfaro et al., 2019). If germination rates decrease with warmer climates, species ranges will likely contract, changing forests composition and favoring invasive species spread in the near future (Hellmann et al., 2008). The timing of germination of dormant seeds requiring chilling and forcing temperatures to germinate is very disturbed by changes in temperatures (Gremer et al., 2020), whereas less is known on how increases in temperature may affect recalcitrant seeds (i.e., desiccation sensitive seeds (Baskin & Baskin, 2001; Obroucheva et al., 2016)), as those of Mediterranean oaks (Amimi et al., 2020), which main requisite to germinate is to avoid seed desiccation rather than reaching a temperature threshold. Both germination rates and timing have a high ecological relevance: germination rates directly influence the regeneration and the contribution of the seeds to the next generation (Baskin & Baskin, 2001; Donohue et al., 2010), whereas the germination timing determines the environment where the plant will develop, and hence its early-stage fitness (Gremer et al., 2020).

Trait co-variation modulates fitness across species ranges (Laughlin et al., 2020), causing compensatory trade-offs at species distribution margins (Willi and Van Buskirk, 2022). Under harsh conditions as those expected at species distribution margins, trade-offs between germination and post-germination traits may define species niches (Donohue et al., 2010). Germination represents the first relationship of the plant with the environment, and it can have a cascade effect on other early-stage fitness-related and functional traits (Donohue et al., 2010; Gremer et al., 2020). For instance, the seed reservoir is mostly used by the cotyledons and hence bigger seeds tend to show faster shoot elongation and first leaf emergence, that can influence the overall seedling growth (Donohue et al., 2010). Likewise, leaf protective pigments as flavonols and anthocyanins can confer protection to stressful environmental conditions as drought and cold temperatures (Chalker-Scott, 1999), in detrimental of other leaf pigments as chlorophyll.

Climate change is favoring the northwards shift of Mediterranean oaks, that are expanding their ranges at their leading edges (Fernández-Manjarrés et al., 2018), becoming natural candidates to increase forest biodiversity in these regions. Among them, *Quercus suber* (cork oak) is a unique forest species with high economic and strong cultural heritage (Ritter and Dauksta, 2013), inhabiting the Maghreb, southern and western Spain, Portugal, east and northeast Italy and southern France, with different gene pools (Magri et al., 2007). Cork oak thick bark (cork) makes it a highly resistant to fire species (Pausas, 2015), a much sought-after feature with the widespread increase of natural fires in the last decades. Furthermore, adult cork oak populations are mostly adapted to drought (Ramírez-Valiente et al., 2010), which confers the species a high resilience to climate change (Pereira et al., 2009). Nevertheless, some populations are declining, mostly due to a combination of drought, pest attacks, fires and management practices (Tiberi et al., 2016). Natural regeneration is often poor, mostly due to rodent and large ungulate herbivores like wild boar predation on the acorns (Herrera, 1995) and grazing management practices, contrasting with the high germination rates observed under laboratory conditions (Amimi et al., 2020; Merouani et al., 2001). Hence, evaluating the real possibilities of cork oak to regenerate under climate change is essential to understand the chances of this species to persist, conquer northern environments and be a good candidate for seed sourcing forest strategies.

Our main goal was to investigate the early stages of cork oak life to evaluate its range-wide vulnerability to climate warming. We sowed 701 acorns from 11 populations covering the entire cork oak distribution range under controlled conditions in climatic chambers emulating the expected spring temperature for 2080. We measured germination and post-germination related traits to i) estimate differences in the timing and probability of populations through Cox’s proportional-hazards models (Cox, 1972); we expected large population differentiation in germination rates and timing (Donohue et al., 2010); ii) investigated the potential adaptation of these traits to spring temperature using transfers using mixed-effects models; we hypothesized that germination and post-germination traits from different populations are adapted to their local temperature as observed in adult cork oaks (Ramírez-Valiente et al., 2009), and that populations with heavier acorns germinate faster and produce faster growing seedlings than lighter ones (Mechergui et al., 2021); iii) predicted the germination and post-germination climatic niches accounting for populations differences and plasticity under current and future conditions (Benito Garzón et al., 2019); we hypothesized that warmer spring temperatures will decrease germination rates and the number of days required to germinate range-wide (Baskin and Baskin, 2001; Donohue et al., 2010), with low germination rates and advanced germination becoming the main barrier for seeding establishment.

## 2. Methods

### 2. 1. Acorn sampling and conservation

During the 2021 autumn, we collected 900 acorns at the same ripening stage (i.e. brown pericarp and without radicle emergence) from 11 populations covering the distribution range of *Quercus suber* (Figure 1). To assure that the maturation point was identical in all populations, we only collected acorns that dropped from trees after shaking their branches. In each population, we collected 10 acorns from 10 different trees (only two at Fuencaliente, Spain, and nine at Padouen, France, in the first case because trees had already dropped all their acorns, and in the second because of the lack of fruiting trees). In total, we collected acorns from 81 mother trees. The individuals sampled were at least 30 meters apart. Coordinates of the populations, mother tree circumference (used as a proxy of maternal effect) and collection date were recorded for all sampled trees (Figure 1). After collection, the acorns were immediately stored in a refrigerator at 1°C for 1 to 3 months, depending on the collection date (Figure 1). The acorns from Tunisia had transportation problems that stopped the cold chain, making acorns germinate before storage and they were discarded from the germination experiment but kept in the chambers for the following post-germination measurements. We immersed acorns in water and floating seeds were removed to select healthy ones before sowing them. In total, 701 acorns were used in the experiment (Figure 1). The experiment started the 08/02/2022 for all the populations and lasted until the seedlings were 28 days old, counted after the first leaves emerged.

**Figure 1.**
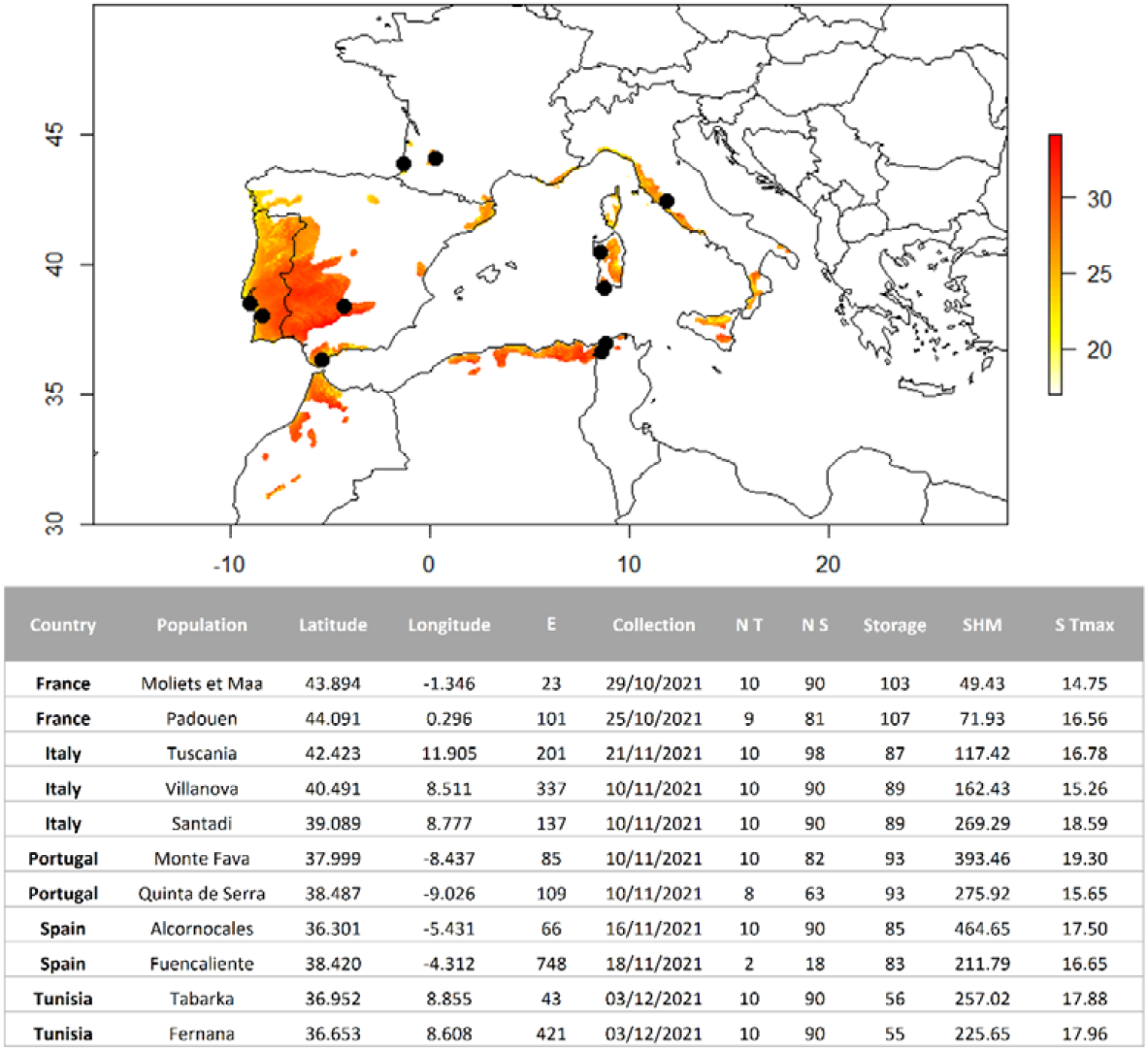
Cork oak acorn origins (map) within the species distribution range from EUFORGEN (Caudullo et al., 2017). Background colors show the spring temperature of 2021 (°C) within the cork oak distribution range. Description of the acorn source populations: E = Elevation (m); NT = number of trees per population; NS = number of acorns per population; Storage = duration of acorn storage at −1°C (days) SHM = Summer Heat Moisture Index averaged for the period of 1901 to 1960; S Tmax = maximum spring temperature averaged for the period of 1901 to 1960.

### 2.2. Experimental design

We used nursery trays (4×5 cells) with a dimension of 6×6 cm with 8 cm depth per cell. In each cell we placed one acorn on a previously added universal substrate (NPK 8-2-7, dry organic matter 80%, conductivity 30mS/m, pH 6.5, water retention 820ml/L). The trays were placed in three climatic chambers (Snijder LABS, micro clima-series) at 15, 20 and 25°C with constant values of 75% relative humidity, 300 μmol.m^-2^.s^-1^ light intensity, and a photoperiod of 13/11 light/dark hours. During the dark phases the temperature was lowered by 5°C in all chambers. Within each shelf the acorns were positioned randomly. We watered the acorns daily with distilled water throughout the experiment to keep optimal humidity for germination.

### 2.3. Phenotypic measurements

We grouped our measurements into germination traits (acorn mass, germination rate and germination timing), and post-germination traits (growth related traits, survival, morphological traits and leaf pigment contents) (Table 1). Acorns were weighted before sowing by groups of then and averaged by each mother tree with a precision of 10^-3^g. After the sowing date, germination was monitored daily during 90 days. The germination time was measured as the number of days between sowing and germination, defined as radical protrusion of at least 2 mm (Table 1). After germination occurred, we measured for each acorn the timing of first leaf emergence. In addition, 28 days after the first leaf emerged, we measured seedling height, number of branches, number of leaves, mean leaf area and leaf pigment contents (chlorophyll content, epidermal flavonoid content, Nitrogen Balance Index (NBI; NBI = Chlorophyll/Flavonoid contents), anthocyanin index). Leaf pigment contents were measured with an optical leaf-clip Dualex (Goulas et al., 2004). We checked the survival status (death/alive) of the seedlings at the end of the experiment and calculated two survival measurements: survival in relation to the number of seeds sown and in relation to the number of seeds germinated (Table 1).

**Table 1.**
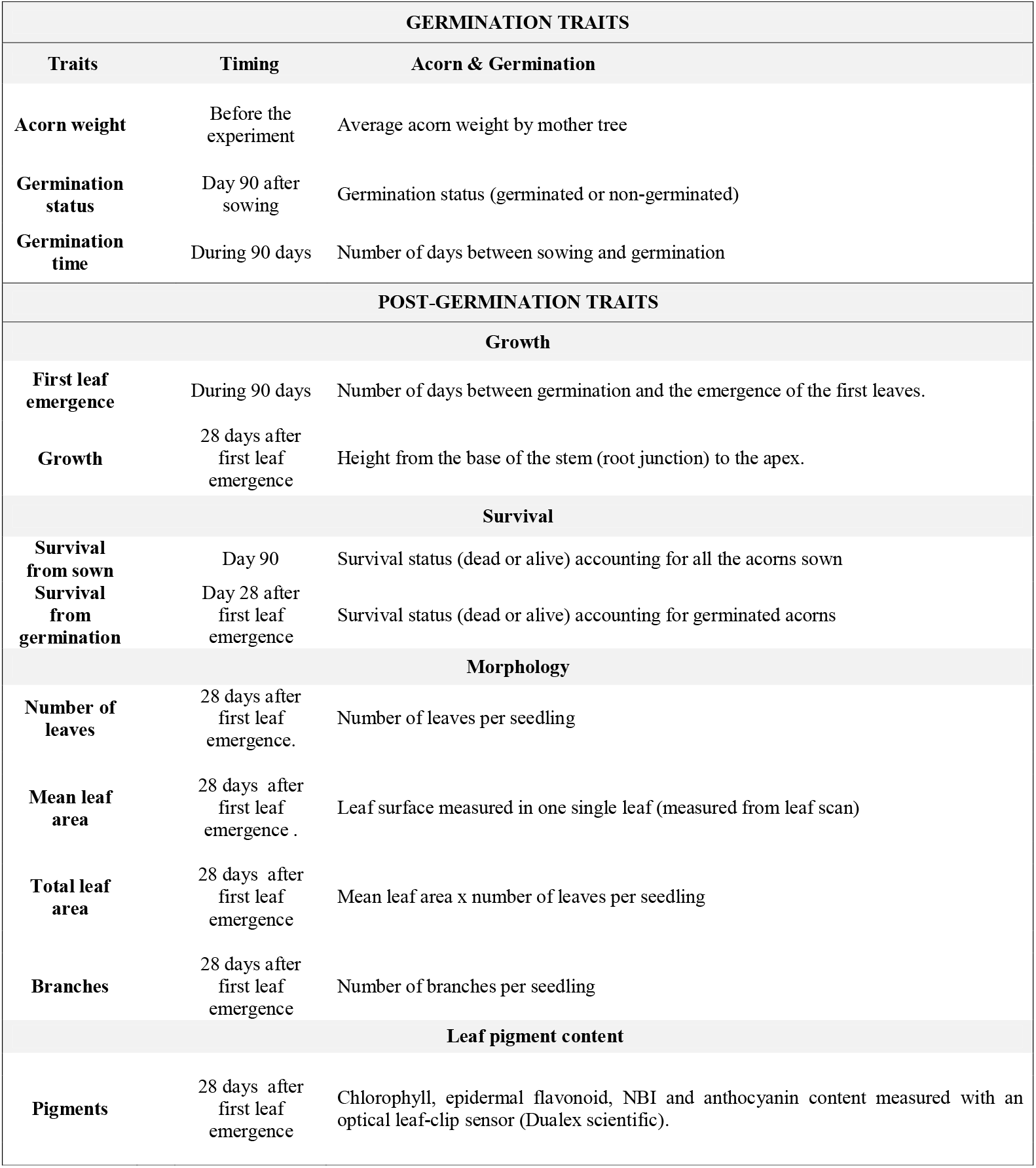
Phenotypic traits measured during the experiment. Germination related traits were measured during 90 days after sowing the acorns. Post-germination traits: growth, morphology and leaf pigment contents were measured 28 days after the first leaf emergence. All the measurements were performed in the climatic chambers except the acorn weight.

### 2.4. Climate data

We used two climatic indexes to identify warm and dry conditions in the models: the Summer Heat Moisture index (SHM) or the Maximum Spring Temperature (MST). The SHM is defined as

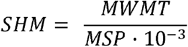

where MWMT is the Mean Warmest Month Temperature (°C) and MSP the Mean Summer Precipitation (mm).

To characterize the adaptation to temperature of the acorns from different origins we calculated a temperature transfer distance for Maximum Spring Temperature (*TD_MST_*) as follows:

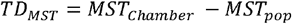

Where *MST_Chamber_* is the diurnal temperature of the climatic chamber that emulates maximum spring temperature expected within the species range under climate change scenarios (15, 20 and 25°C) and *MST_pop_* is the maximum spring temperature of the population averaged for the period of 1901 to 1960, with the rationale that it was the climate that shaped adaptation of mother trees to the local conditions. Positive transfer distances indicated that the temperature of the climatic chamber was warmer than the population spring temperature.

To predict the germination and post-germination climatic niches, we extracted the SHM and MST values and their corresponding raster maps for current and 2080 RCP 8.5 conditions from the ClimateDT (https://www.ibbr.cnr.it//climate-dt/). (Supplementary Figure S1).

## 3. Calculation

### 3.1. Cox proportional-hazards model of germination over time

We used the Cox’s proportional-hazards models to test for the temperature treatment and the differences in germination timing among populations. The proportional-hazards function *h*(*t*|*x*) that estimates the failure rate at time t. *h*(*t*|*x*) at time *t* depends on the covariates *x*_i_ = (*x*_i1_,....., *x*_in_) and it takes the form:

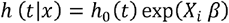

where *x*_i_ represents the populations sown in each chamber, *β* the coefficient for the base of the exponential factor, *h*_0_(*t*) is the baseline hazards and *t* is the germination over time. The exp(*β*) represents the hazard ratios (HR). In our case, a HR represents the risk for an acorn to no germinate between time 0 and t. The main assumption of Cox’s proportional-hazards model is that the hazards between any groups are proportional over time.

We added two covariates, the temperature of the climatic chamber and a quadratic term for temperature to simulate the optimum temperature curve expected for germination (Watt and Bloomberg, 2012). Models were fitted with the *survival* R package (Therneau, 2020).

### 3.2. Mixed-effects models of trait population responses to spring temperature transfer distances and maternal effects

To understand local adaptation of populations to spring temperature we built independent mixed-effects models of germination and post-germination traits as a function of the maximum spring temperature transfer distance *TD_MST_*. Models took the following general form:

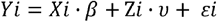

Where *Yi* is the trait measured for each acorn/seedling *i, Xi* · *β* is the fixed block, *Zi* · *ν* is the random block and *ε* is the error.

In particular, we used generalized logistic mixed-effects models with a natural logit link function (GLMM; glmm R package) for the binomial variables germination status, survival from sown, and survival after germination; and linear mixed-effects (LME; lme4 R package; (Bates et al., 2018)) for all the other traits with a Gaussian distribution. Models took the following specific form:

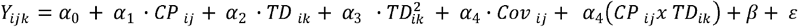

where *Y_ijk_* = trait response of the *i*th acorn of the *j*th population in the *k*th climatic chamber; *Cov* = trait covariate of the *i*th acorn in the *k*th climatic chamber; *CP* = climate at the population site of the *i*th acorn of the *j*th population; *TD* = climatic transfer distance at the climatic chamber of the *i*th acorn of the *j*th population in the *k*th chamber; β = random effects and ε = residuals. The model included the interaction of the *TD* and *CP.*

The fixed-effects included *CP, TD, TD*^2^, *Cov* and the interaction between *CP* and TD. Where *CP* = MST or SHM, depending on the model information criterion (AIC) (Akaike, 1973); TD = *TD_MST_*; *Cov* included the storage time (number of days that the acorns were storage at 1°C), the circumference at breast height of the mother tree (as a maternal effect) and the average acorn mass per mother tree. The random-effects included the population and mother tree nested to the population.

The variance explained by the models was measured by marginal (R^2^M) and conditional (R^2^C) variances, i.e. by only considering fixed effects or fixed and random effects together, respectively (function r.*squaredGLMM,* package *MuMIn* (Barton 2015)). Finally, the capacity of generalization of the model was assessed by the R Pearson coefficient comparing the observed versus predicted values using a cross validation method with independent data (64% of the data used for calibration and the remaining 36% for validation). All the variables were standardized subtracting the mean value and divided by one standard deviation before running the models.

### 3.3. Fitness related traits spatial predictions under current and future spring temperatures

We used the mixed-effects models to perform trait-based niche predictions for those fitness-related traits with statistically significant population terms (Benito Garzón et al., 2019). The predictions were performed under current and future conditions (2080; RCP 8.5) and they were exclusively based on the fixed-effects of the models. Covariates were fixed to the average per population value. The predictive models were restricted to the current cork oak distribution range (Caudullo et al., 2017). Raster images were analyzed using the *raster, sp* and *gdal* libraries of R (R Development Core Team, 2022).

## 4. Results

### 4.1. Treatment and population effects on the probability of germination over time

The global germination rate was 20.83% (16.94% at 15°C, 27.54% at 20°C and 17.90% at 25°C), meaning that only 146 acorns germinated from a total of 701 sown. The timing of germination showed great differences depending on the population and the temperature treatment (Supplementary Figure S2). Our model follows proportional hazards (Supplementary Table 1). Both germination timing and temperature -linear and quadratic effects- were significant to explain the probability of germination for all the populations except Quinta de Serra which germination rate was independent of germination timing (Table 2). The differences in temperature treatments and its quadratic effect were both significant (Table 2) and negative.

**Table 2.** Cox’s proportional-hazards model parameters. N = number of acorns sown by climatic chamber; N events = number of germinated acorns; β = coefficient of regression; exp(β) is the coefficient for the base of the exponential factor; z is the z value of the model; p is the p-value and 95%CI are the confidence intervals; Temperature = temperature of the treatment. Temperature^2^ = quadratic effect of the temperature. Significant p-values (p< 0.05) are shown in bold.

| | $\beta$ | $\exp(\beta)$ | z | p | 95% CI |
| --- | --- | --- | --- | --- | --- |
| **Alcornocales** | -1.03 | 0.27 | -2.36 | **0.018** | 0.10-0.80 |
| **Monte Fava** | -1.20 | 0.30 | -2.13 | **0.033** | 0.10-0.91 |
| **Padouen** | -1.49 | 0.23 | -2.57 | **0.010** | 0.07-0.70 |
| Quinta da Serra | -0.40 | 0.67 | -0.72 | 0.473 | 0.22-2.02 |
| **Santadi** | -1.47 | 0.16 | -2.57 | **0.002** | 0.05-0.51 |
| **Moliets et Maa** | -2.32 | 0.10 | -3.67 | **<0.001** | 0.03-0.34 |
| **Tuscania** | -1.85 | 0.16 | -3.19 | **0.001** | 0.05-0.49 |
| **Villanova** | -1.52 | 0.22 | -2.66 | **0.008** | 0.07-0.67 |
| **Temperature** | -0.86 | 0.42 | -3.01 | **0.003** | 0.24-0.74 |
| **Temperature<sup>2</sup>** | -0.02 | 1.02 | 3.10 | **0.002** | 1.01-1.04 |
| N = 701 |  |  |  |  |  |
| N events = 146 |  |  |  |  |  |

The maximum difference among populations in number of days to reach 50% germination estimated by the Cox’s proportional-hazards model was 32 days occurred at 20°C for acorns coming from Fuencaliente (36 days) and those coming from Moliets et Maa (68 days). The difference to reach 50% of germination between these two extreme populations was 30 days at 25°C and 28 days at 15°C (Supplementary Table S2; Figure 4).

The highest probability to germinate over time was predicted for the Fuencaliente and Quinta de Serra populations, and the lowest for the Moliets et Maa one, regardless the temperature treatment (Figure 2).

**Figure 2.**
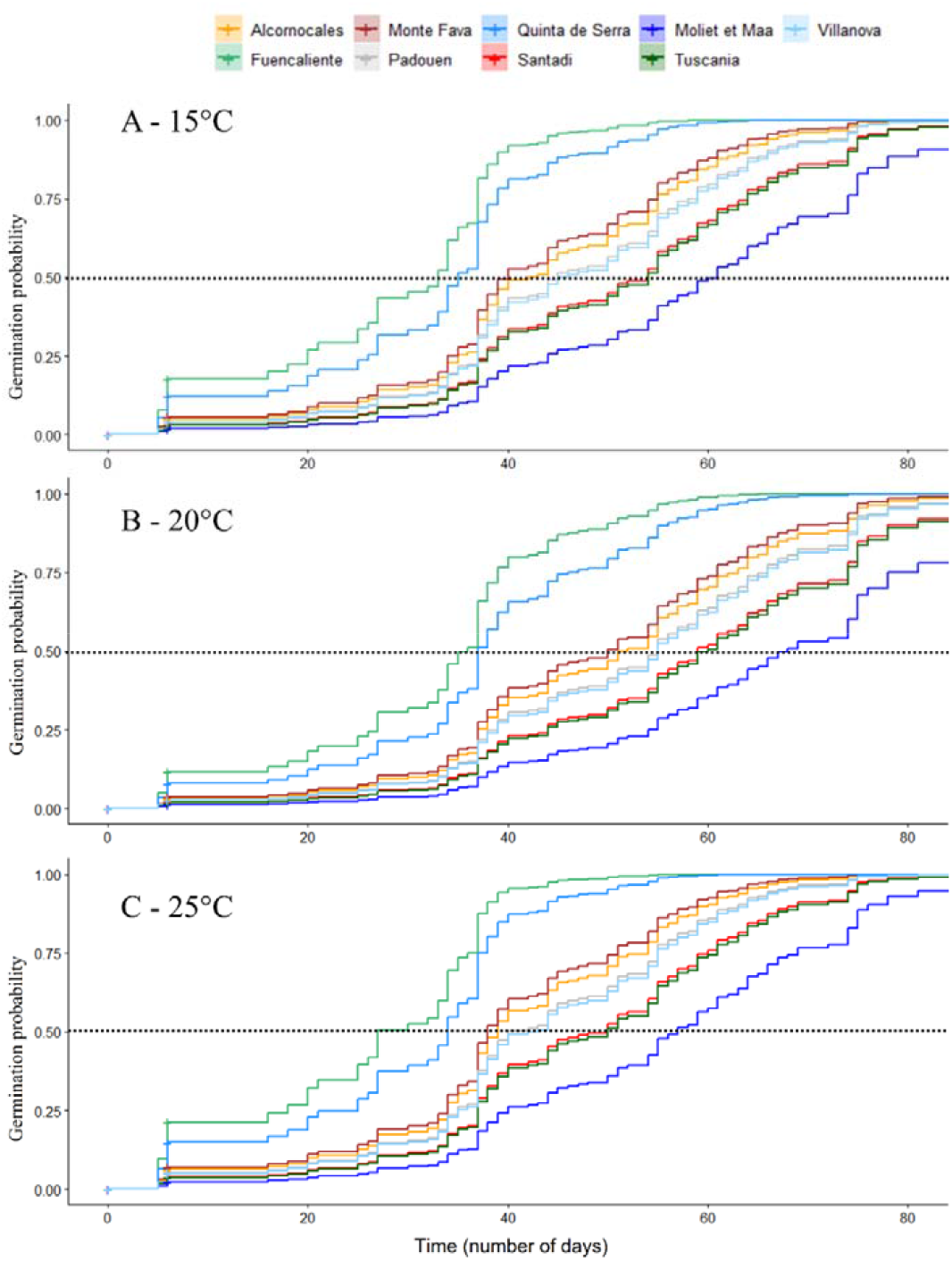
Probability population germination curves over time estimated by the Cox proportional hazards models (Table 3) for 15° (A), 20° (B) and 25°C (C). The dash line indicates the 50% probability of germination.

### 4.2. Mother circumference, storage duration and acorn mass effects on germination and post-germination traits

The circumference of the mother tree and the summer heat moisture index at the source of the population had significant and positive effects explaining acorn mass (Table 3). The model had high performance (Table 3). Large mother trees from drier and warmer populations (Monte Fava, Alcornocales and Santadi) yielded heavier acorns than small trees, whereas wetter and colder populations as Moliets et Maa did not show any differences in their acorn mass related to the mother tree size (Supplementary Figure S3). In all the other models we removed the covariate circumference of the mother tree as it was correlated with the acorn mass.

**Table 3.** Fixed and random effects of linear mixed-effects models of germination and postgermination traits measured in the climatic chambers. All the traits are measured after germination, except the survival from sown that includes non-germinated acorns. All traits were analyzed with linear mixed-effect models except survival and probability of germination ones that used binomial linear mixed-effects models. Acorn mass model did not include the random effects. I= Estimate= coefficient of each variable within the model; R^2^M= marginal variance, attributed to fixed effects; R^2^C= conditional variance, including fixed and random effects; R-squared = total variance for acorn mass linear model; Pearson R = Pearson coefficient for observed versus predicted values with independent data (jackknife validation after 100 bootstrap); PC = Population Climate; MST = Maximum Spring Temperature of the populations averaged for the period of 1901 – 1960; *TD_MST_* = *MST_Chamber_* – *MST_pop_*; SHM = Summer Heat Moisture index of the populations averaged for the period of 1901 – 1960; Circum = circumference of the mother tree. NBI = Nitrogen Balance Index; MLA = Mean Leaf Area. t values over 2 and under −2 represent a significant estimate. Significant estimates with p <0.01 are marked in bold.

|  |  |  | Fixed- Effects |  |  |  |  |  | Random-<br>mother tree/population | Effects | Model performance |  |  |
| --- | --- | --- | --- | --- | --- | --- | --- | --- | --- | --- | --- | --- | --- |
| Trait | Intercept<br>Estimate;<br>t-value | PC (SHM<br>or MST)<br>Estimate; t<br>value | Covariates |  |  |  | TD<sub>MST</sub><sup>2</sup><br>Estimate; t<br>value | PC x covariate<br>Estimate ; t<br>value | Variance | stddev | R<sup>2</sup>M | R<sup>2</sup>C | R Pearson |
|  |  |  | Acorn<br>mass<br>Estimate; t<br>value | Storage time<br>Estimate; t<br>value | Circum<br>Estimate; t<br>value | TD<sub>MST</sub><br>Estimate<br>; t value |  |  |  |  |  |  |  |
| Germination traits |  |  |  |  |  |  |  |  |  |  |  |  |  |
| Acorn mass | 5.63; **99.60** | SHM: 0.93;<br>**14.77** |  |  | 0.68; **11.54** | - | - | 0.46; **8.09** | - | - | 0.40 | - | 0.63 |
| Germination<br>probability | -2.07; -1.30 | MST: 0.04;<br>0.38 | 0.05; 0.78 | -0.01; -0.66 | - | - | -0.02; **-2.98** | 0.01; **2.65** | 0.59 | 0.77 | 0.03 | 0.18 | 0.44 |
| Germination timing | 59.30;<br>**10.03** | SHM: -<br>0.04; -1.74 | -0.94; -1.18 | 0.15; 0.31 | - | - | -0.22; **-3.55** | 0.003; **2.03** | 19.82 | 4.45 | 0.10 | 0.30 | 0.59 |
| Post-germination traits |  |  |  |  |  |  |  |  |  |  |  |  |  |
| Growth |  |  |  |  |  |  |  |  |  |  |  |  |  |
| Growth Day 28 | 9.15; **-5.88** | MST:<br>1.049; 1.75 | 0.72; **2.12** | -0.03; -0.63 | - | - | 0.12;<br>**4.07** | 0.03; **4.09** | 2.39 | 1.55 | 0.30 | 0.54 | 0.81 |
| Time to first leaves<br>emergence | 158.70;<br>**4.53** | MST: -<br>6.52; **-3.21** | 0.03; 0.92 | 0.02; 1.71 | - | - | -0.08; -0.5 | -0.17; **-3.38** | 0.00 | 0.00 | 0.34 | 0.45 | 0.74 |
| Survival |  |  |  |  |  |  |  |  |  |  |  |  |  |
| Survival from sown | -8.15; **-3.20** | MST:<br>0.35; **2.42** | 0.28; 1.86 | 0.05; 0.66 | - | 1.35;<br>**2.48** | -0.03; **2.43** | -0.07; **-2.44** | 0.99 | 0.99 | 0.07 | 0.29 | 0.39 |
| Survival of germinated<br>acorns | -4.72; -1.54 | MST: 0.30;<br>1.66 | 0.03; 0.42 | -0.02; -0.56 | - | - | -0.02; 0.28 | -0.06; 0.16 | 0.13 | 0.35 | 0.09 | 0.12 | 0.35 |
| Morphology |  |  |  |  |  |  |  |  |  |  |  |  |  |
| Number of leaves | 9.39; 4.46 | SHM: 0.00;<br>0.43 | 1.32; **3.74** | 0.07; 0.71 | - | - | 0.06; 1.26 | 0.02; 0.09 | 1.87 | 0.37 | 0.32 | 0.44 | 0.37 |
| MLA | -0.42; 0.06 | MST: 0.56;<br>1.34 | 0.47; **2.68** | -0.01; -0.12 | - | - | -0.06; **-2.11** | 0.01; 0.58 | 2.02 | 1.42 | 0.14 | 0.32 | 0.72 |
| Total MLA | 5.82; 0.04 | MST: 5.51;<br>0.56 | 17.36; **5.37** | -0.45; -0.66 | - | - | -0.40; 0.62 | 0.14; 1.06 | 207.8 | 14.42 | 0.06 | 0.06 | 0.25 |
| Ramification | 0.05; 0.35 | MST: 0.05;<br>0.54 | 0.03; 0.90 | 0.001; 0.25 | - | - | 1.29; 0.69 | 0.00; -0.26 | 0.03 | 0.17 | 0.01 | 0.07 | 0.36 |
| Leaf pigment content |  |  |  |  |  |  |  |  |  |  |  |  |  |
| NBI | 26.88; **8.65** | SHM: -<br>0.00; -0.95 | -1.00; **-2.45** | 0.06; 0.26 | - | - | 0.05; 1.11 | 0.01; **3.21** | 13.61 | 3.69 | 0.24 | 0.45 | 0.69 |
| Chlorophyll | -8.16; -0.67 | MST: 2.12;<br>**3.00** | -0.51; -1.79 | -0.07; -0.77 | - | - | -0.13; **-2.88** | 0.14; **7.50** | 0.63 | 0.79 | 0.55 | 0.56 | 0.76 |
| Flavonols | 1.05; 13.14 | SHM: 0.00;<br>**2.12** | 0.01; 1.32 | <0.001; -1.47 | - | - | -0.00; -0.93 | 0.000; 1.78 | 0 | 0.08 | 0.23 | 0.42 | 0.69 |
| Anthocyanins | 0.00; -0.25 | SHM:<br>0,1.97 | <0.001;<br>**3.36** | <0.001; 0.33 | - | - | -0.00; -0.36 | -0.00, -0.21 | 0.00 | 0.01 | 0.09 | 0.19 | 0.58 |

The storage duration did not have a significant effect in any of the germination and post-germination traits evaluated (Table 3). Acorn mass had a significant effect in the following post-germination traits at day 28 after first leaf emergence (Table 3): growth, number of leaves, MLA, total MLA, NBI and anthocyanin models.

### 4.3. Effects of spring temperature transfer distances on germination and post-germination traits

The germination probability model showed a negative quadratic significant relationship between the germination probability and the *TD_MST_*, indicating the existence of an optimum where the germination probability is the highest and that decreased towards both sides of transfer distances. The interaction between the MST and the *TD_MST_* was significant, indicating a genetic x environment interaction (Table 3). The co-variates storage time and acorn mass were not significant in the germination model. All populations showed their optimum germination probability in warmer than their local climate conditions (TD_MST_ > 0). Positive TD_MST_ always predicted that populations from warmer spring origins (Monte Fava, Santadi, Alcornocales) would had higher germination probabilities than those from colder spring origins (Padouen, Quinta de Serra, Villanova, Moliets et Maa) (Figure 3B).

**Figure 3.**
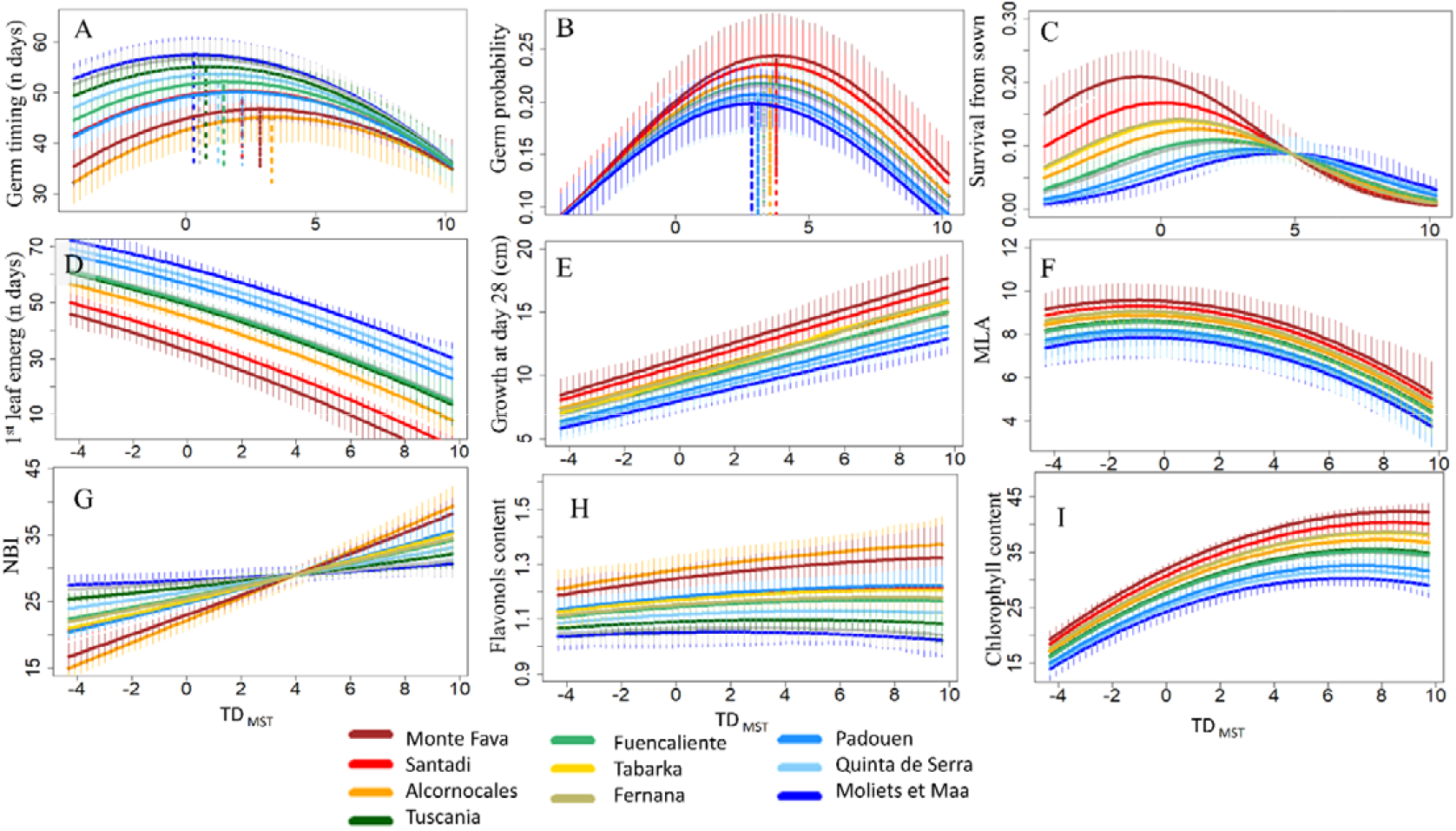
Population responses of germination (excluding acorn mass) and post-germination traits to maximum spring temperature transfer distances (TD_MST_). Transfer distances indicate the differences in temperature between the climatic chambers and the maximum spring temperature of the populations. Dotted vertical lines show the differences in highest (optimum) values for germination probability and timing along TD_MST_. Vertical segments represent confident intervals after 100 simulations. Germination probability, germination timing, survival from sown and first leaf emergence was not considered for Tunisian populations (i.e. Tabarka and Fernana) because they were already germinated when they arrived to the laboratory. MLA = mean leaf area; NBI = Nitrogen Balance Index.

The negative quadratic significant relationship of the germination timing with the *TD_MST_* indicated the existence of an optimum number of days for germination. The significant interaction term between SHM and *TD_MST_* indicated the existence of genetic environmental interaction (Table 3). The co-variates storage time and acorn mass were not significant in the germination time model. The number of days to germinate decreased along the SHM of the populations origin, i.e. the populations coming from dry environments (Alcornocales, Monte Fava, Santadis) germinated earlier than those from a less dry origin and this happens between -*4*<*TD_MST_*<9 (Moliets et Maa, Padouan, Tuscania) (Figure 3A).

The number of days required after germination to first leaf emergence decreased significantly along spring temperature (Table 3; MST estimate = −6.52). The interaction between the MST of the population and the *TD_MST_* was also significant, indicating an environment x genetic significant effect (Table 3). Acorns from colder populations (Moliets et Maa, Villanova, Quinta de Serra, Padouen) required more days until first leaves emerged than those coming from warmer origins (Monte Fava, Santadi, Alcornocales) (Figure 3D). Furthermore, less number of days for first leaves emergence was needed when sowing acorns in warmer sites than their local origin.

The interaction between population spring temperature and TD_MST_ and the acorn mass were the only significant drivers for seedling growth 28 days after leaf emergence (Table 3). Populations from warmer origins grew more when planted in warmer conditions than populations from colder origins (Figure 3E).

From both survival measurements, only the survival from sown model had significant terms (Table 3) and hence was the only that we used as survival model hereafter. The MST of the population, the transfer distances (*TD_MST_* and 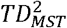) and their interaction were significant in the model (Table 3). Populations from warmer origins survived more when sown under local conditions (*TD_MST_*= 0; Figure 3C), whereas populations from cooler origins survived more when planted under warmer conditions (*TD_MST_* > 0; Figure 3) and populations from a milder origin survived more when planted in slightly warmer than origin conditions (approx. 0< *TD_MST_*<5 Figure 3C).

The number of leaves model depended exclusively on the acorn mass (Estimate = 1.32; t-value = 3.74; R^2^M = 0.32; R^2^C = 0.44: R Pearson = 0.37). Mean leaf area (MLA) had two significant terms, specifically, the acorn mass and the quadratic *TD_MST_* (Table 3). Seedlings coming from warmer origins had a larger MLA than those from colder origins. All populations tended to present the highest MLA under local conditions (i.e. at *TD_MST_* = 0; Figure 3F). The Total Mean Leaf Area unique significant term was acorn mass (Estimate = 17.36; t-value = 5.37).

Nitrogen Balance Index (NBI), chlorophyll and flavonols showed significant model terms (Table 3). Seedlings from drier populations showed higher NBI than those from wetter populations when planted at warmer than origin temperatures (positive transfer distance), whereas the pattern shifted when NBI was evaluated for negative transfer distances (Figure 3G). Seedlings from warm and dry origins showed higher chlorophyll and flavonols contents than wetter and colder populations, respectively. For all populations, both pigments increased when seedlings grew under warmer conditions than their origins (Figure 3H and 3I).

### 4.4. Spatial predictions of fitness-related traits

We only performed trait-niche predictions for those traits with at least one significant climatic variable in the mixed-effects models(Table 3): germination probability, germination timing, growth at day 28 after first leaf emergence (hereafter growth), and survival from sown (hereafter survival). Please note that models are exclusively based on the fixed-effects, i.e. the relationship of the trait with *TD_MST_*, SHM or MST of the population origin, depending on the model.

The probability of germination under current and 2080 climate scenario was very similar within the cork oak distribution (germination differences between 2080 and 2021 ranged from −0.08 to 0.04; Figure 4A), only slightly higher in southern Portugal and North Africa (Supplementary Figure S4).

**Figure 4.**
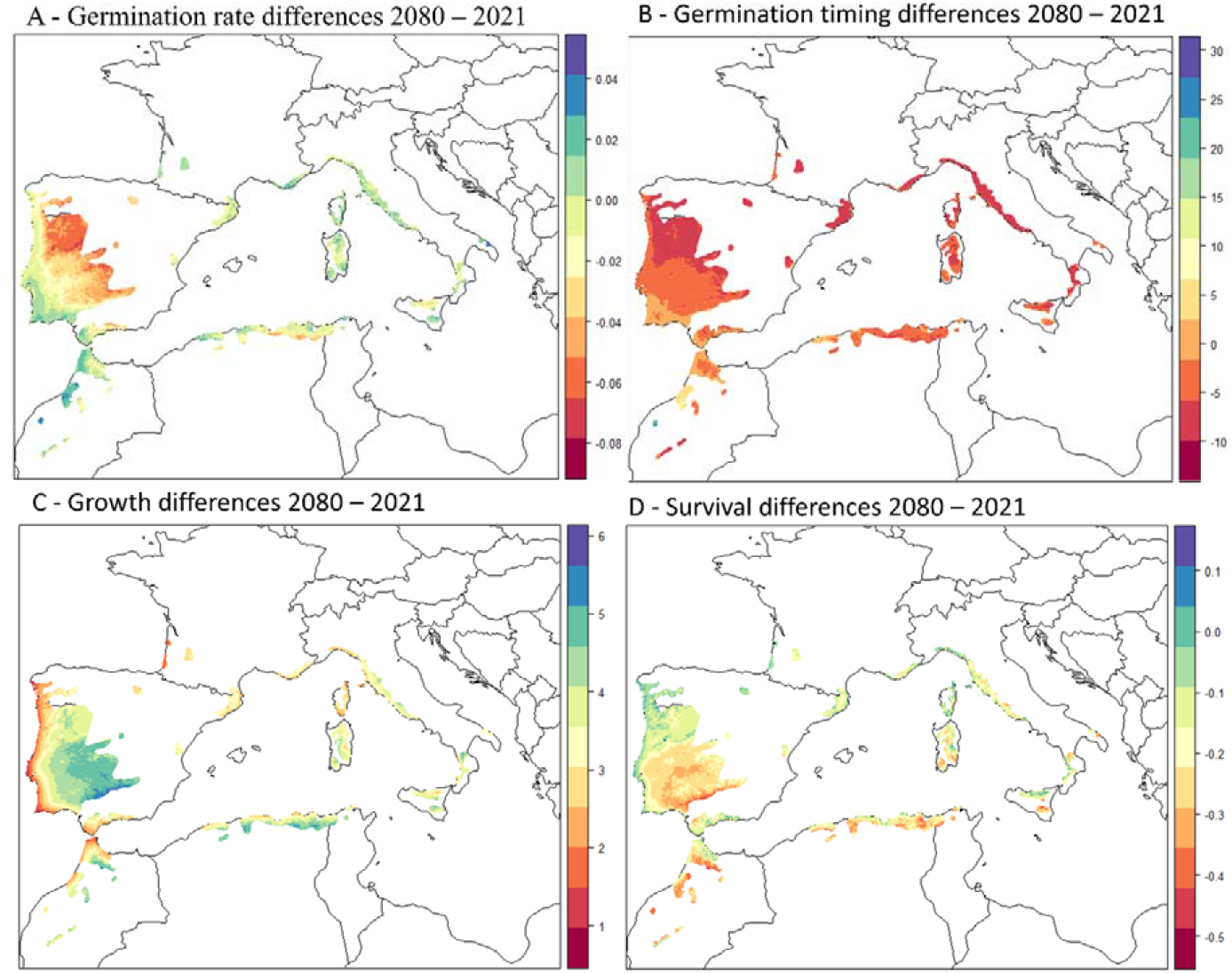
Differences in the trait-based niche predictions for current and 2080 RCP 8.5. predicted from mixed-effects models. Top left: germination probability; Top right: germination time (number of days); Bottom left: growth at day 28 after first leaf emergence (mm); Bottom right: probability of survival.

The differences among populations in the number of days required for germination ranged from more than 60 days in the Nouvelle Aquitaine (south-western France) to less than 30 in some parts of Morocco (Figure 4; Supplementary Figure S4). Differences in predictions for the 2080 - 2021 period showed an overall decrease in the number of days required for germination, with northern populations predicted to germinate 12 days before than today (Figure 4B).

The current probability of survival was predicted to be slightly lower in the northern part of cork oak distribution range than in the southern part. Under the 2080 RCP 8.5 scenario survival was predicted to decrease all over its distribution range, with a large decrease in central Spain and eastern Portugal (Figure 4C; Supplementary Figure S4).

Predicted differences in seedling growth for 2080 were very small (ranging between 1 and 6 mm). RCP 8.5 2080 spring temperature conditions were predicted to favour seedling growth across the entire cork oak range, with higher growth predicted for central Spain and eastern Portugal populations (Figure 4D).

## 5. Discussion

### 5.1. Storage effect and desiccation filtering

Mediterranean oaks acorn capacity to keep their germination potential from the autumn drop until spring depends mostly on the humidity that they receive during winter, making seed desiccation sensitivity a key functional trait that strongly influences seed germination and seedling recruitment (Baskin and Baskin, 2001; Obroucheva et al., 2016). In our experiment, the desiccation suffered by cork oak acorns during the storage has probably selected those acorns more tolerant to desiccation, reducing the expected germination rate when compared with other experiments under controlled conditions (from the ~90% observed by Amimi et al., 2020; Merouani et al., 2001 to the ~21% that we observed)..

### 5.2. Maternal and acorn mass effects on germination, survival, growth and pigment contents

Our findings showed that larger mothers produce heavier acorns under drier conditions, which likely reflects the differences in allocation resources across developmental stages (Roach and Wulff, 1987). Young (small) trees mostly allocate their resources for growth, whereas older (big) trees allocate more resources for reproduction than for growth. Regardless the mother circumference, populations from colder origins (notably the two at the leading edge of cork oak distribution, Moliets et Maa and Padouen) always had small acorns (Supplementary Figure S3), which may indicate a tolerance to cold trade-off in cold leading-edge populations (Ramírez-Valiente et al., 2009).

The lack of relationship between seed mass and germination probability that we found is common in plants, as the seed reserve depot is mostly used by the cotyledons, and hence seed mass is generally more related to seedling growth than to germination (Obroucheva et al., 2016). We expected to find a positive relationship between germination timing and acorn mass, as embryos from larger seeds require more time to develop and hydrate before tegument rupture than those from small seeds (Norden et al., 2009), but the timing of germination only depended on the population origin climate and the climatic transfer distance (Table 3).

As expected (Donohue et al., 2010; Mechergui et al., 2021; Merouani et al., 2001), acorn reserves allocation was mostly used by the cotyledons and hence had a positive effect in growth. The positive relation of acorn mass with leaf anthocyanin content and the negative one with the leaf NBI content (i.e. chlorophyll/flavonols) may indicate a higher allocation of resources to drought protection pigments as anthocyanin and flavonols than to chlorophyll production (Chalker-Scott, 1999).

### 5.3. Large differences in germination timing among populations are triggered by spring temperature

In most plants, germination timing is under strong natural selection (Cendán et al., 2013). In the case of recalcitrant species, which main barrier to germination is desiccation (Obroucheva et al., 2016), we could expect that adaptation to warm conditions may have led to population differentiation (Donohue et al., 2010), as shown by our results (Figure 2 and Figure 3A). More importantly, the lower number of days required by warm populations to germinate (Figure 3A; Figure 4B and Supplementary Table S3), is likely an adaptive response to avoid early drought in warmer regions (Vizcaíno-Palomar et al., 2014), and might also be an adaptation to irregular spring precipitation that often occurs in the Mediterranean basin. However, it is difficult to determine the advantages of an earlier versus a delayed germination for a plant. Early germination avoid competition from siblings and drought stress happening later in the season, whereas delayed germination could benefit seedling survival by avoiding other unfavorable environmental conditions (Donohue et al., 2010), as for instance late cold events as those that occur in the northernmost part of cork oak distribution range. Furthermore, advanced germination timing related to warmer springs is important for assisted population programs, which attempt to move populations from warmer origins to colder parts of a species distribution range to compensate for climate change (Aitken and Bemmels, 2016).

### 5.4. Evidences of mal-adaptation in germination probability and survival

Locally adapted populations reach their optimum trait values at home climate (i.e. TD = 0) (Rehfeldt et al., 2002), hence when optimum trait values are found towards lower or higher than zero transfer distances, it can be interpreted as a population mal-adaptation. We found mal-adaptation in germination probability for all populations considered, as they germinated more at higher temperatures than those found at their local origins (optimum germination occurs at TD_MST_ > 0) (Figure 3). This positive TD_MST_ suggests that germination rates could slightly increase until warming exceeds approximately 4°C, point at which germination may decrease as shown by the spatial predictions (Figure 4A). In the case of survival, we found similar mal-adaptation patterns as those found for adult tree growth (Fréjaville et al., 2020), with populations from cold origin adapted to warmer-than-home conditions (TDMST >0). The mal-adaptation of these populations towards warmer environments may favor northern populations survival under warmer climates as shown by the survival spatial predictions (Figure 4D). Although this mal-adaptation towards warming climates might be encouraging results for a species that is being proposed as alternative in restoration programs under climate change (Vessella & Schirone, 2022), they must be taken with caution, as the final establishment of seedlings in the field may be affected by other factors such as predation, competition, drought, and extreme cold events that may reduce the final chances of cork oak regeneration.

### 5.5. Germination and post-germination niches under climate change

Our trait-based niche models point out the south-western Iberian and northern Morocco as the most vulnerable regions for the establishment of cork oak in the future. This is in agreement with the results based on landscape genomics, that highlight a strong disruption of gene-environment relationship in these regions (Vanhove et al., 2021). However, the differences in survival and growth predictions between current and future spring temperature remain very small, and changes in germination rates due to increase spring temperatures in the future are negligible (Figure 4). In comparison, the large advance in germination predicted by the models all across the cork oak range would be decisive for the establishment of the species in the near future. For instance, early germination in the northern margin of the cork oak distribution range may imply an increased risk of cold events that can be deleterious for young seedlings (Amimi et al., 2020), but it can also increase seedlings chances to survive by avoiding excessive temperatures linked to climate change and allowing the extension of the growing season before the onset of the dry Mediterranean summer.

## 6. Conclusions

Germination timing is the most sensitive to climate early stage trait for cork oak. This is not surprising, as phenology events are very disrupted by climate change (Visser and Both, 2005). However, whereas the advanced fruiting and flowering phenology is well documented (Ma et al., 2022), germination timing is an understudied trait that ultimate determines tree regeneration and is very disrupted by climate change. We claim that more studies on germination timing are urgently needed to understand trees vulnerability to climate.

The choice of climate-resilient species in restoration, reforestation and assisted migration programs is generating an intense debate because it depends on multiple ecological, economic and social factors (Findlater et al., 2022). Cork oak ecological characteristics as resistance to fire and adaptation to drought, as well as it economic importance for cork extraction, makes it a natural candidate for such programs in temperate Europe. Our results showing the earlier germination related to warmer temperatures is essential for seed sourcing strategies (Aitken and Bemmels, 2016), that need to consider the balance between planting fast growing populations and the risk of cold events related to early germination that may hamper the natural regeneration of the translocated material.

## Abbreviations

SHM: Summer Heat Moisture index
MST: Maximum Spring Temperature
MWMT: Mean Warmest Month Temperature
MSP: Mean Summer Precipitation
*TD_MST_*: Climatic transfer distance for MST
R^2^M: Marginal variance estimated by mixed-effects models. It includes the variance related to the fixed effects.
R^2^C: Conditional variance estimated by mixed-effects models. It includes the variance related to mixed and random effects.
NBI: Nitrogen Balance Index. It measures the ratio between clorophyll and flavonoid leaf pigments.

## Acknowledgements

NVP was supported by IJC2020-044557-I. All the seeds were collected following the The Nagoya Protocol on access to genetic resources and the fair and equitable sharing of benefits arising from their utilization to the convention on biological diversity. We are very grateful to all our colleagues, friends and family that helped us with acorn collection. We thank Julien Goullier from the ‘Le Liège Gascon’ association for showing us the best Nouvelle Aquitaine cork oak populations and for his help with acorn collection. We also thank Pedro Ruiz, Abel González y María Angeles Pacheco, the technical team of ‘Finca La Almoraima’ (Cadiz, Spain) for their support and help collecting acorns. Andrea Bertoldi, Amelia and Olivia Bertoldi Benito and David Smith enthusiasm and help was essential to collect the French acorns; Amit Mansukhani and Enya Vizcaino Mansukhani help was essential to collect the Spanish ones.

## Notes

### Competing Interest Statement

The authors have declared no competing interest.

