## Supplementary Material for "Germination timing under climate change: warmer springs favor early germination of range-wide cork oak populations"

|  | **χ^2^** | **df** | **p** |
| --- | --- | --- | --- |
| Population | 7.54 | 8 | 0.48 |
| Temperature | 1.38 | 1 | 0.24 |
| Temperature^2^ | 1.31 | 1 | 0.25 |
| GLOBAL | 8.83 | 10 | 0.55 |

**Supplementary Table S1**. χ^2^ test of proportional hazards in the Cox’s proportional-hazards model. Non-significant values (p-values > 0.05) indicate that we reject the hypothesis of non-proportionally, and hence that our model follows proportional hazards. df: degree of freedom

| **Population** | **15°C** | | | | **20°C** | | | | **25°C** | | | |
| --- | --- | --- | --- | --- | --- | --- | --- | --- | --- | --- | --- | --- |
|  | **N** | **NG** | **ND** | **CI** | **N** | **G** | **ND** | **CI** | **N** | **NG** | **ND** | **CI** |
| **Alcornocales** | 30 | 5 | 43 | 37-56 | 30 | 14 | 51 | 44-61 | 30 | 9 | 39 | 37-51 |
| **Fuencaliente** | 6 | 1 | **33** | 21-NA | 6 | 3 | **36** | 27-NA | **6** | 0 | **27** | 20-NA |
| **Monte Fava** | 28 | 6 | 40 | 37-55 | 28 | 8 | 50 | 40-61 | 26 | 5 | 38 | 37-52 |
| **Padouen** | 27 | 3 | 45 | 38-61 | 27 | 8 | 55 | 45-67 | 27 | 4 | 40 | 37-59 |
| **Quinta de Serra** | 22 | 6 | 35 | 32-40 | 23 | 8 | 37 | 35-50 | 18 | 3 | 34 | 27-39 |
| **Santadi** | 30 | 5 | 54 | 43-69 | 30 | 6 | 59 | 54-75 | 30 | 6 | 50 | 39-63 |
| **Moliets et Maa** | 30 | 5 | **61** | 54-78 | 30 | 2 | **68** | 59-NA | 30 | 4 | **57** | 46-81 |
| **Tuscania** | 33 | 2 | 54 | 44-72 | 32 | 9 | 60 | 54-75 | 32 | 6 | 50 | 39-63 |
| **Villanova** | 30 | 7 | 45 | 39-59 | 30 | 7 | 55 | 46-67 | 30 | 4 | 43 | 37-58 |
| **Total number of sown acorns** | 236 |  |  |  | 236 |  |  |  | 229 |  |  |  |
| **Total Germination rate (%)** | 16.94 |  |  |  | 27.54 |  |  |  | 17.90 |  |  |  |
| **Average number of days to reach 50% germination** |  |  | 45.56 |  |  |  | 52.33 |  |  |  | 42.00 |  |

**Supplementary Table S2**. Estimation of the number of days needed for each population to reach 50% germination using the Cox’s proportional-hazards models in the three temperature treatments. N: number of acorns sown; NG: number of acorns germinated; ND: number of days needed to reach 50% germination; CI: confidence interval at 95%. Numbers in bold remark the minimum and maximum number of days to reach 50% germination in each treatment.

| **Population** | **Temperature = 27°C** | | **Temperature = 30°C** | |
| --- | --- | --- | --- | --- |
|  | **Ndays** | **95%CI** | **Ndays** | **95%CI** |
| **Alcornocales** | 37 | 33-50 | 25 | 6-NA |
| **Fuencaliente** | 21 | 6-NA | 6 | 5-NA |
| **Monte Fava** | 35 | 30-51 | 21 | 6-NA |
| **Padouen** | 37 | 34-64 | 26 | 6-NA |
| **Quinta de Serra** | 27 | 18-44 | 6 | 5-NA |
| **Santadi** | 38 | 36-61 | 30 | 18-NA |
| **Moliets et Maa** | 46 | 37-NA | 35 | 25-NA |
| **Tuscania** | 39 | 37-63 | 32 | 18-NA |
| **Villanova** | 37 | 34-59 | 27 | 6 -NA |
| **Average number of days to reach 50% germination** | 35.22 |  | 23.11 |  |

**Supplementary Table S3**. Estimation of the average number of days to reach 50% germination using Cox proportional-hazards model (Table 3) for the temperatures of 27° and 30°C.

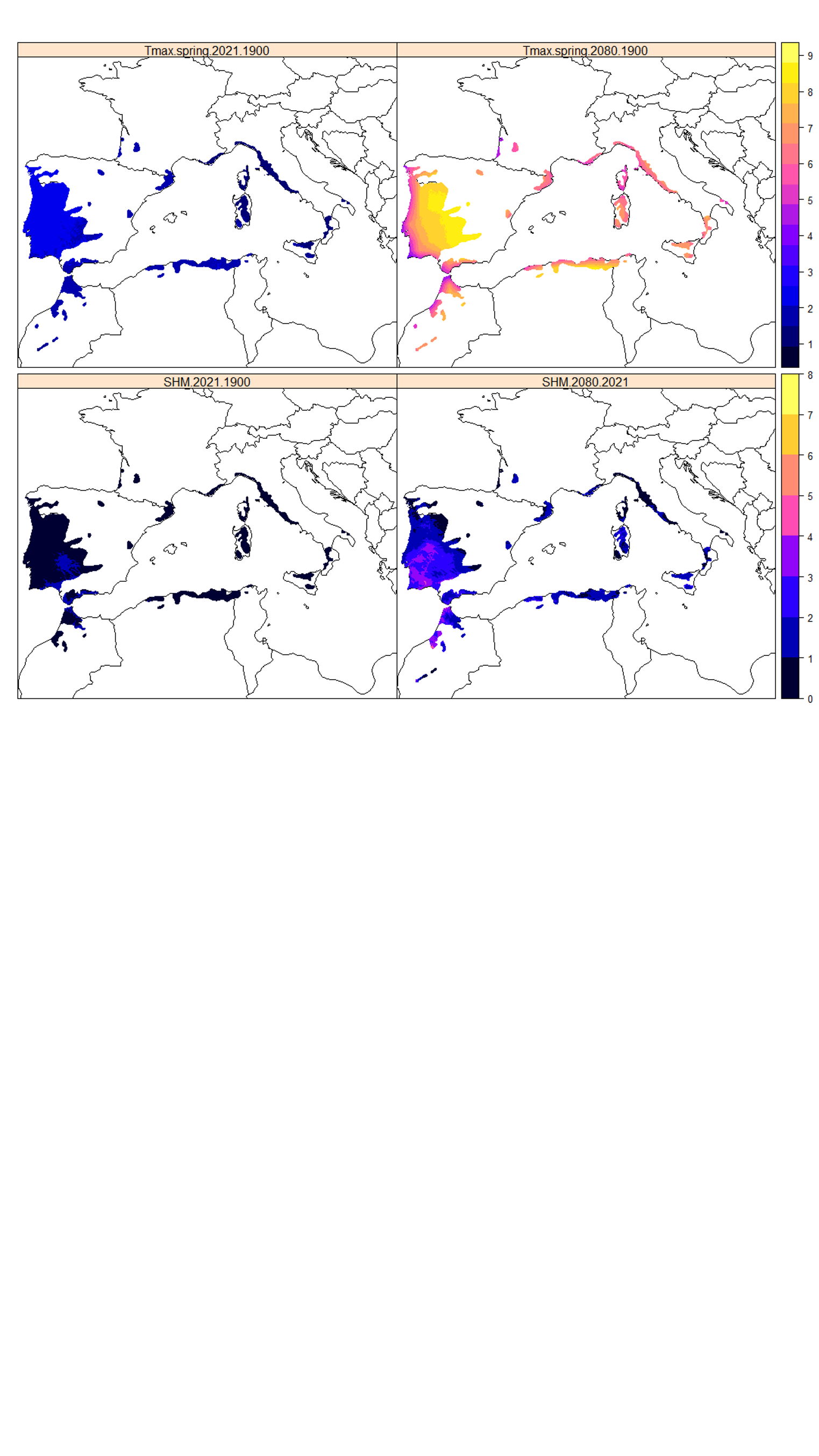

**Supplementary Figure S1**. Top: Differences in maximum spring temperature between 2021 and 1900 (left) and 2080 (RCP 8.5) and 1900 (right) as used in the models for calculating the transfer distances, and differences in SHM between 2080 (RCP 8.5) and 2021 as used in the models. Maps are restricted to the cork oak distribution range.

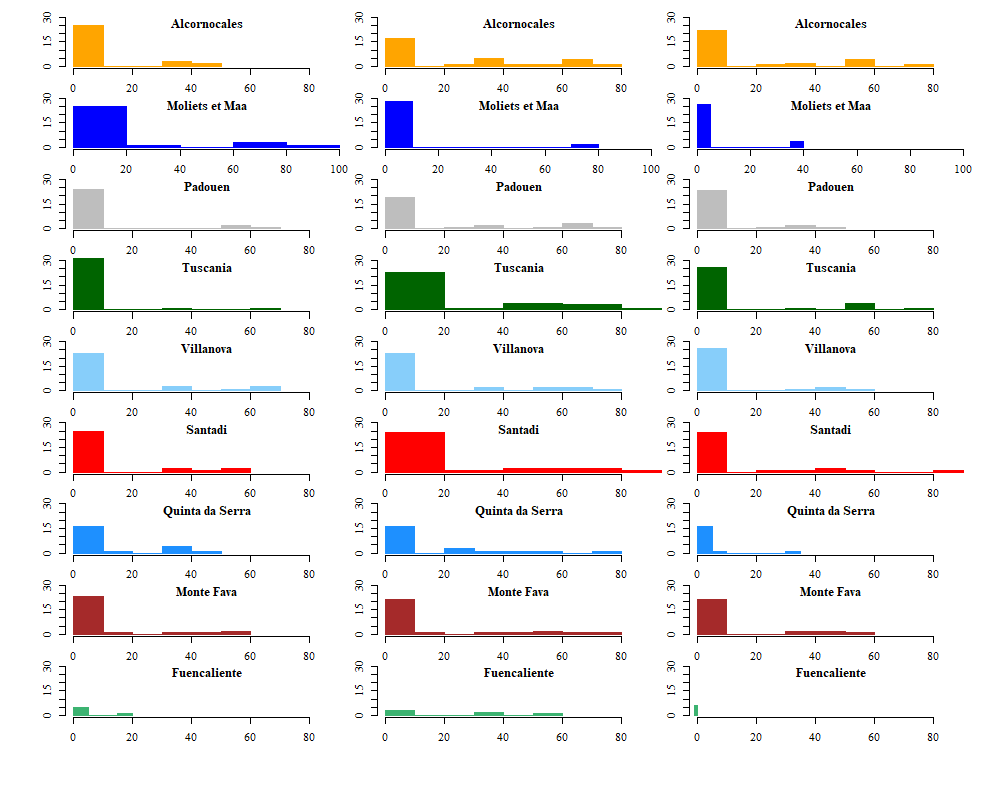

**Supplementary Figure S2** – Number of seeds germinated over time per population as observed in the three climatic chambers (left 15°C, center 20°C, and right 25°C). x axis = number of days required for germination; y axis = number of germinated seeds.

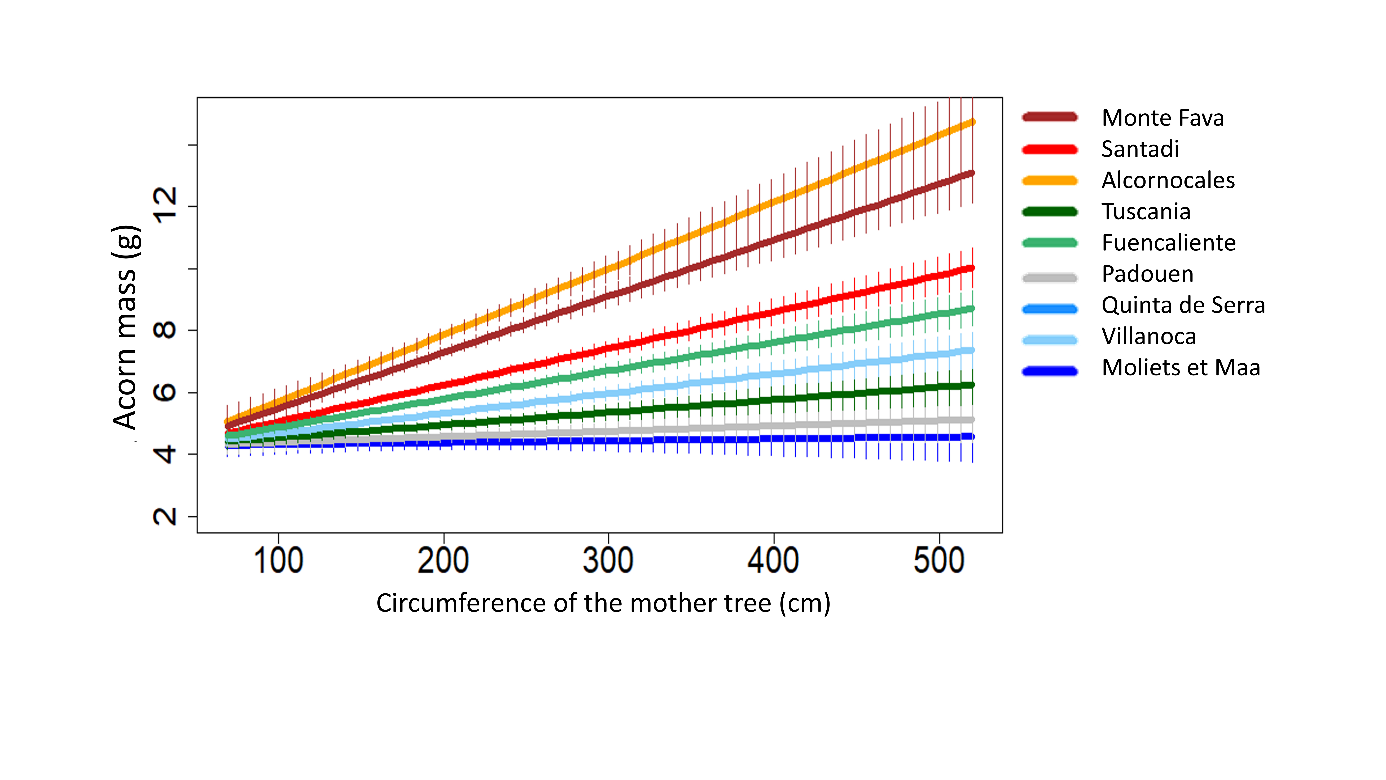

**Supplementary Figure S3** – Population responses of acorn mass to the circumference of the mother trees. Vertical segments represent confident intervals after 100 simulations. Acorns from Tunisian populations (i.e. Tabarka and Fernana) were not considered as they were already germinated when arrived to the laboratory.

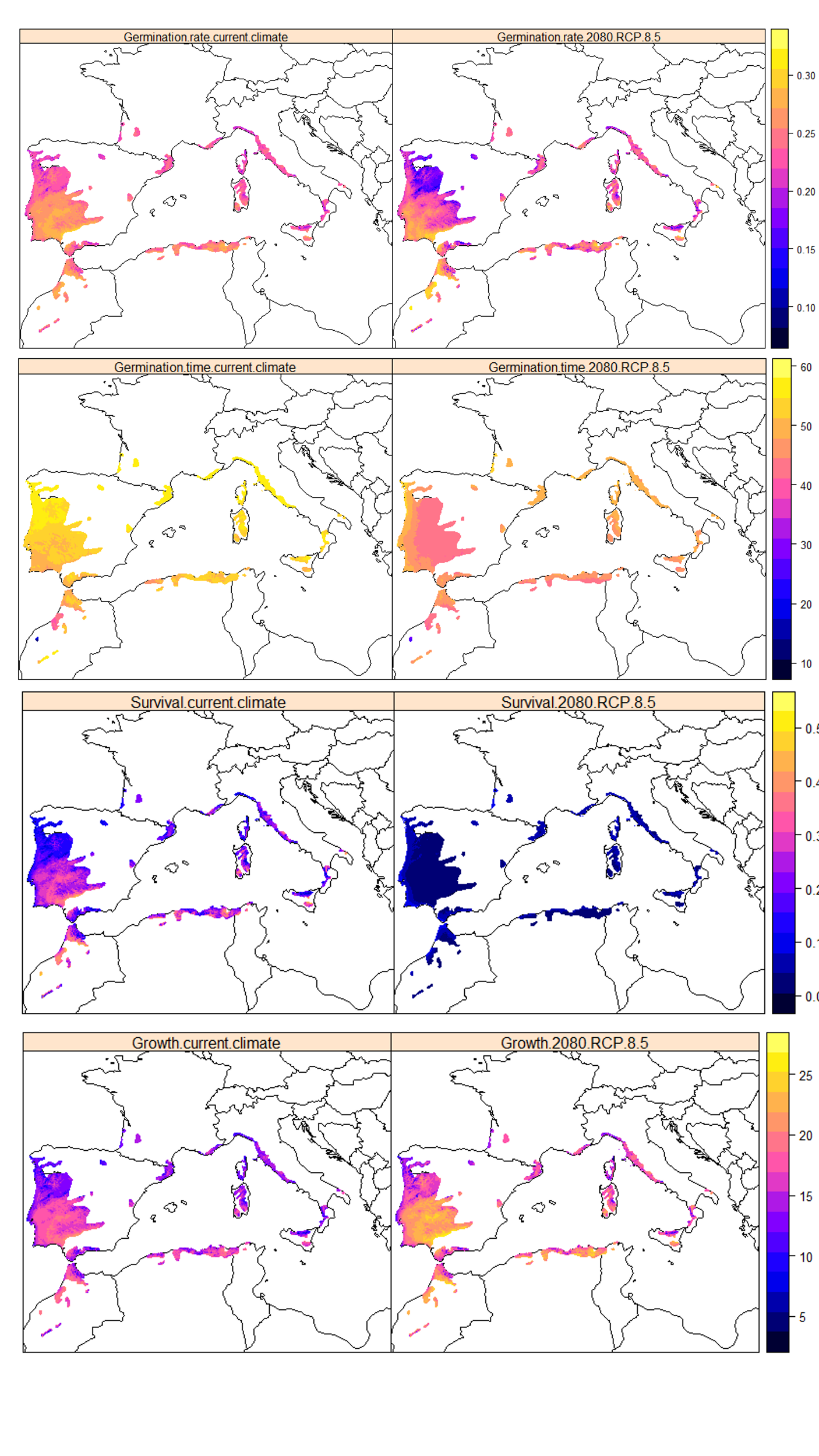

**Supplementary Figure S4.** Spatial predictions of fitness-related traits (germination rate, germination timing, survival and growth) under current (baseline 2011 – 2021; left) and future climate conditions (2080 RCP 8.5; right).
